# Evidence that disruption of apicoplast protein import in malaria parasites evades delayed-death growth inhibition

**DOI:** 10.1101/422618

**Authors:** Michael J. Boucher, Ellen Yeh

## Abstract

Malaria parasites (*Plasmodium* spp.) contain a nonphotosynthetic plastid organelle called the apicoplast, which houses essential metabolic pathways and is required throughout the parasite life cycle. Hundreds of proteins are imported across 4 membranes into the apicoplast to support its function and biogenesis. The machinery that mediates this import process is distinct from proteins in the human host and may serve as ideal drug targets. However, a significant concern is whether inhibition of apicoplast protein import will result in a “delayed-death” phenotype that limits clinical use, as observed for inhibitors of apicoplast housekeeping pathways. To assess the growth inhibition kinetics of disrupting apicoplast protein import, we targeted a murine dihydrofolate reductase (mDHFR) domain, which is stabilized by the compound WR99210, to the apicoplast to enable inducible blocking of apicoplast-localized protein translocons. We show that stabilization of this apicoplast-targeted mDHFR disrupts parasite growth within a single lytic cycle in an apicoplast-specific manner. Consistent with inhibition of apicoplast protein import, stabilization of this fusion protein disrupted transit peptide processing of endogenous apicoplast proteins and caused defects in apicoplast biogenesis. These results indicate that disruption of apicoplast protein import avoids delayed-death growth inhibition and that target-based approaches to develop inhibitors of import machinery may yield viable next-generation antimalarials.

**Importance:** Malaria is a major cause of global childhood mortality. To sustain progress in disease control made in the last decade, new antimalarial therapies are needed to combat emerging drug resistance. Malaria parasites contain a relict chloroplast called the apicoplast, which harbors new targets for drug discovery, including import machinery that transports hundreds of critical proteins into the apicoplast. Unfortunately, some drugs targeting apicoplast pathways show delayed growth inhibition, which results in a slow onset-of-action that precludes their use as fast-acting, frontline therapies. We used a chemical biology approach to disrupt apicoplast protein import and showed that chemical disruption of this pathway avoids delayed growth inhibition. Our finding indicates that prioritization of proteins involved in apicoplast protein import for target-based drug discovery efforts may aid in the development of novel fast-acting antimalarials.

## Introduction

*Plasmodium* spp. parasites cause malaria and are responsible for over 200 million human infections and over 400,000 deaths annually (1). Despite a reduction in malaria-related mortality in the past 15 years, emerging resistance to frontline antimalarials necessitates the continued development of new chemotherapies (2, 3). One key source of drug targets is the apicoplast, a nonphotosynthetic plastid organelle found in many apicomplexan pathogens (4, 5). The apicoplast produces essential metabolites required for parasite replication (6). Apicoplast function requires the import of over 300 nuclear-encoded gene products into the organelle, including biosynthetic enzymes and pathways that support organelle biogenesis and maintenance (7). Most apicoplast proteins are 1) synthesized on cytosolic ribosomes, 2) trafficked to the apicoplast via the endoplasmic reticulum (ER), and 3) translocated across the 4 apicoplast membranes. This multistep import pathway involves more than a dozen proteins, including homologs of the translocation and ubiquitylation machinery typically involved in ER-associated degradation (ERAD) (8-12) as well as homologs of the TOC and TIC machinery found in plant plastids (9, 13-15).

Apicoplast protein import machinery are potential drug targets, but a key unresolved question is whether inhibition of apicoplast protein import causes a “delayed-death” phenotype that has been observed for inhibitors of apicoplast housekeeping functions (16, 17). During delayed death, parasite growth is unaffected during the first lytic cycle of inhibitor treatment but is severely inhibited in the second lytic cycle even after drug removal. This *in vitro* phenotype manifests as a slow onset-of-action that limits clinical use of these drugs. Unfortunately, there are no inhibitors known to act directly on apicoplast protein import, precluding direct assessment of growth inhibition kinetics. Furthermore, most genetic tools available in *Plasmodium* parasites act at the DNA or RNA levels (18), which can result in different growth inhibition kinetics than direct chemical inhibition of that same target (19-21). Destabilization domains that conditionally target proteins for degradation by the cytosolic ubiquitin-proteasome enable protein-level disruption (22, 23), but these systems are not suitable to study apicoplast-localized proteins, which are inaccessible to the cytosolic proteasome.

Given these limitations, we developed a protein-level tool to dissect growth inhibition kinetics following disruption of apicoplast protein import. A murine dihydrofolate reductase (mDHFR) domain which can be conditionally stabilized by a small molecule has been used to characterize protein translocation steps during yeast mitochondrial protein import *in vitro* (24) and, more recently, during export of malarial proteins across the parasitophorous vacuole (PV) membrane into the host red blood cell (25, 26). Strikingly, in the context of malarial protein export, mDHFR stabilized with the compound WR99210 can block the translocon to prevent export of other *Plasmodium* proteins and disrupt parasite growth (27). Given that there are at least three putative translocation steps during apicoplast protein import (7), we reasoned that we could use a similar strategy to block apicoplast translocons and disrupt protein import in a manner that resembles inhibition with a small molecule (Fig. 1A).

**Figure 1.**
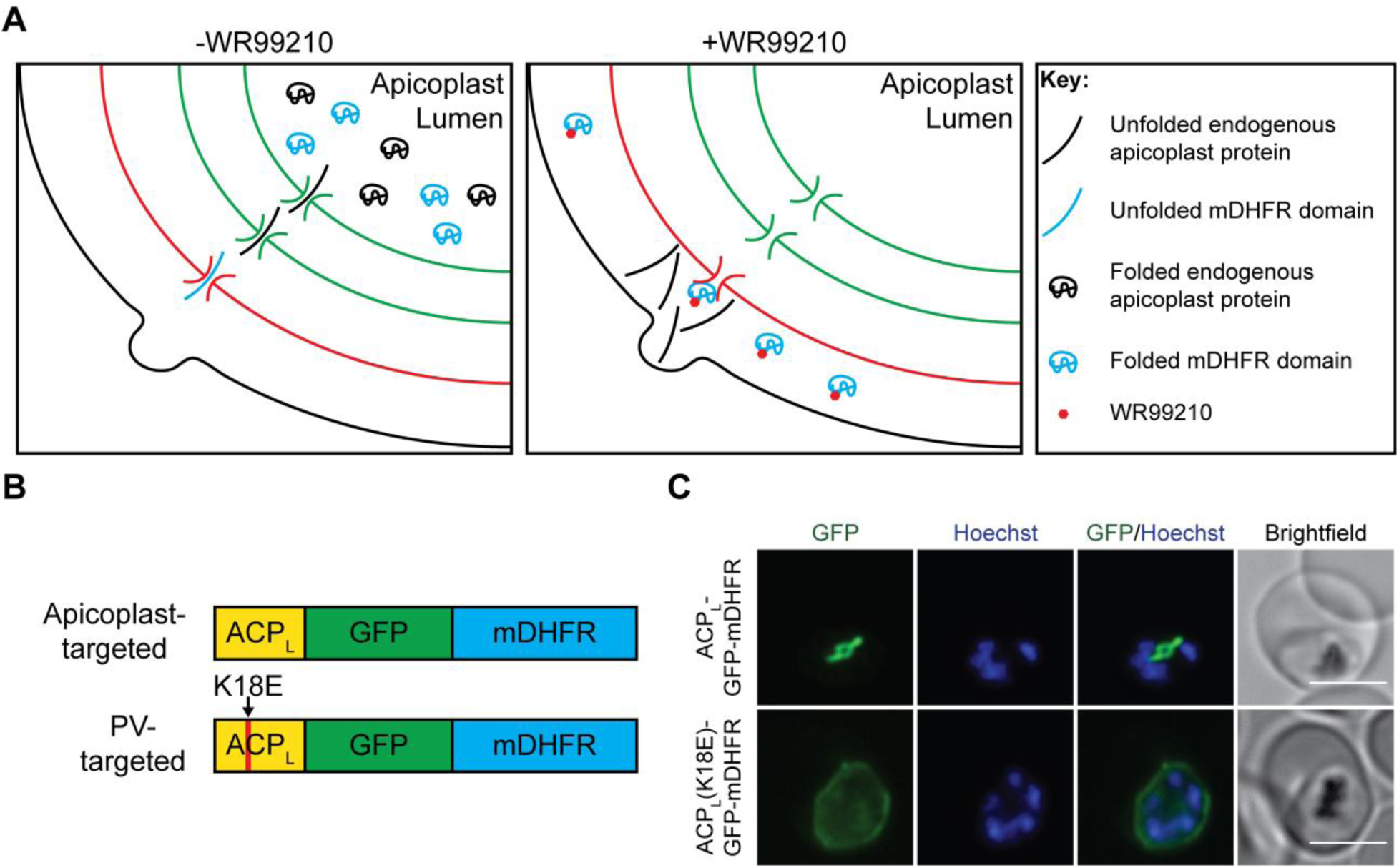
Generation of a protein-level conditional tool to disrupt apicoplast protein import. (A) Model for protein-level disruption of apicoplast protein import by a conditional stabilization domain. In the absence of WR99210, an apicoplast-targeted mDHFR domain successfully targets to the apicoplast lumen along with endogenous apicoplast proteins. Addition of WR99210 stabilizes the mDHFR domain to prevent unfolding, which causes the domain to stall during translocation across one or multiple apicoplast membranes and block import of essential endogenous apicoplast cargo. (B) Schematic (not to scale) of constructs targeting GFP-mDHFR to the apicoplast via the ACP leader sequence or to the PV via a mutant ACP leader sequence. (C) Live-cell imaging of GFP-mDHFR fusions. Nuclei were stained with Hoechst. Brightness/contrast adjustments were not held constant between the two cell lines due to differences in GFP fluorescence intensity in the apicoplast versus the PV. Scale bars, 5 µm.

## Results and Discussion

To generate a protein-level conditional tool for disruption of apicoplast protein import, we targeted a GFP-mDHFR fusion to the apicoplast via the *N*-terminal leader sequence of acyl carrier protein (ACP) (Fig. 1B). As a negative control, we generated a cell line expressing the same fusion with a lysine 18 to glutamate (K18E) mutation in the ACP leader sequence that renders this transit peptide nonfunctional and causes mistargeting to the PV (28). Both GFP-mDHFR fusions were expressed in *P. falciparum* Dd2^attB^ parasites (29) and localized to the expected compartments (Fig. 1C).

To test whether stabilization of these GFP-mDHFR fusions disrupted parasite growth, we treated ring-stage parasites with increasing doses of WR99210 and assessed parasitemia after 3 days as a read-out for parasite growth inhibition during the first lytic cycle. Parental Dd2^attB^ parasites, which express a WR99210-resistant human DHFR allele, were unaffected at the WR99210 concentrations tested (Fig. 2A). Parasites expressing the PV-localized ACP_L_(K18E)-GFP-mDHFR fusion from the *attB* site were also insensitive to WR99210. In contrast, parasites expressing the apicoplast-targeted ACPL-GFP-mDHFR fusion showed dose-dependent growth inhibition in response to WR99210 (Fig. 2A).

**Figure 2.**
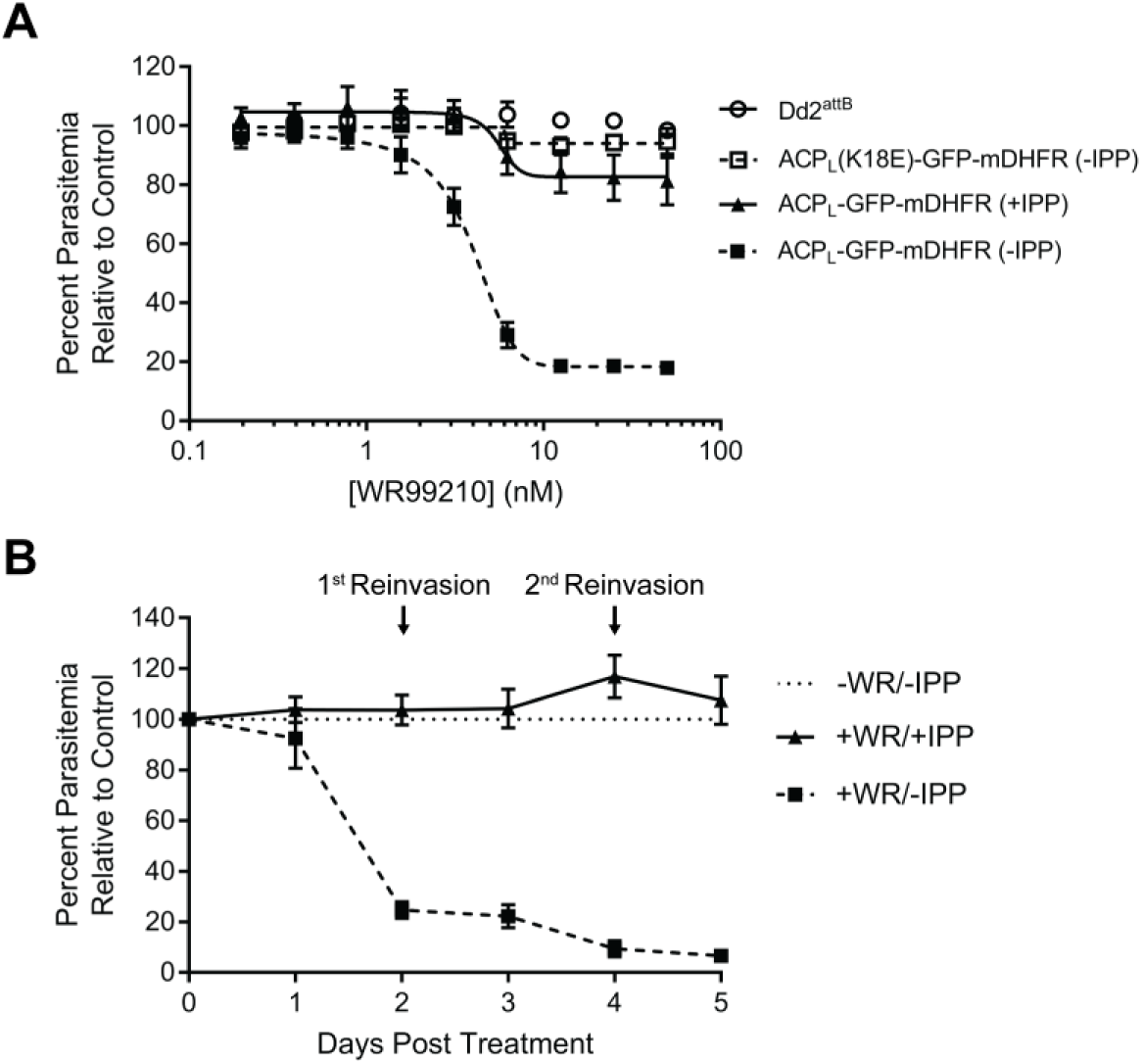
Conditional stabilization of apicoplast-targeted GFP-mDHFR causes apicoplast-specific growth inhibition in the first lytic cycle. (A) Growth of parental Dd2^attB^, ACP_L_-GFP-mDHFR, and ACP_L_(K18E)-GFP-mDHFR parasites after 3 days in response to increasing doses of WR99210. ACP_L_-GFP-mDHFR parasites were assayed in both the presence and absence of 200 µM IPP. (B) Growth of ACP_L_-GFP-mDHFR parasites in the presence of 10 nM WR99210 over a 5-day time course. Parasites were grown in the presence or absence of 200 µM IPP, and parasitemia was normalized to an untreated control at each time point. Error bars in both panels represent standard deviation of the mean of 3 biological replicates. Biological replicates in (A) were performed in technical triplicate.

During *in vitro* culture of blood-stage *P. falciparum*, biosynthesis of the isoprenoid precursor isopentenyl pyrophosphate (IPP) is the only essential function of the apicoplast. As such, supplementation with exogenous IPP can rescue apicoplast defects and can be used to identify apicoplast-specific phenotypes (30). IPP supplementation reversed the WR99210 sensitivity of parasites expressing ACP_L_-GFP-mDHFR in both a 3-day dose-response experiment and over a 5-day time course (Fig. 2A and B). These results indicate that the WR99210 sensitivity conferred by ACPL-GFP-mDHFR is due to disruption of an apicoplast-specific pathway.

Given our hypothesized mechanism-of-action inhibiting apicoplast protein import (Fig. 1A), we expected that ACP_L_-GFP-mDHFR stabilization would cause global disruption of the apicoplast, including organelle maintenance and biogenesis, leading to apicoplast loss. To assess the status of the apicoplast, we assayed the presence of the apicoplast genome in WR99210-treated, IPP-rescued parasites by quantitative PCR (qPCR). WR99210 treatment caused a decrease in the apicoplast:nuclear genome ratio beginning at 1 day post-treatment and near-complete loss of the apicoplast genome after 2 lytic cycles (Fig. 3). The loss of the apicoplast genome, a key marker of the apicoplast, indicates that chemical stabilization of ACP_L_-GFP-mDHFR-expressing parasites caused apicoplast loss.

**Figure 3.**
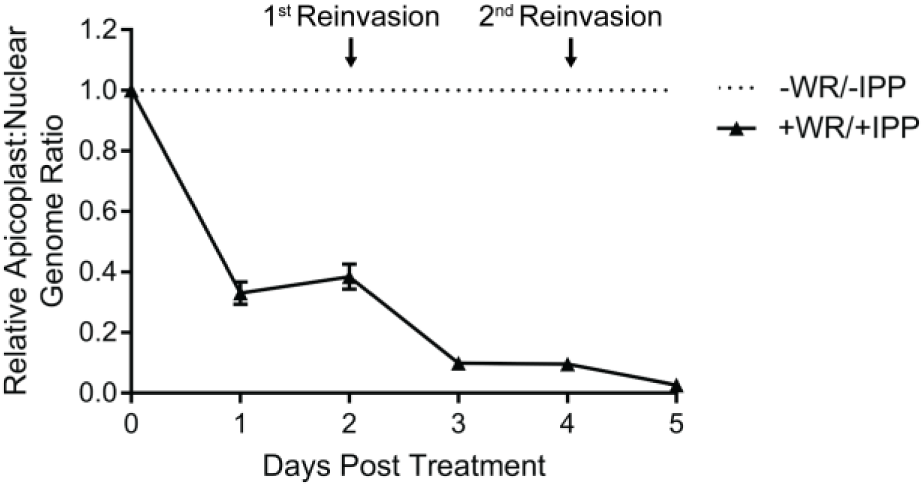
Conditional stabilization of apicoplast-targeted GFP-mDHFR causes an apicoplast biogenesis defect. The relative apicoplast:nuclear genome ratio of ACP_L_-GFP-mDHFR parasites grown in 10 nM WR99210 and 200 µM IPP was measured by qPCR over 5 days of treatment. Values are normalized to an untreated control at each time point. Error bars represent standard deviation of the mean of 3 biological replicates, each analyzed in technical triplicate.

Sorting of apicoplast cargo occurs in the parasite endomembrane system, after which proteins are thought to traffic to the apicoplast by vesicle transport. While we expected that stabilization of ACP_L_-GFP-mDHFR would disrupt transit of apicoplast cargo across the apicoplast-localized ERAD-and TOC/TIC translocons, there was a possibility that this stabilized fusion protein could disrupt the upstream sorting process and prevent apicoplast proteins from ever reaching the organelle. Previous data suggest that defects in sorting apicoplast cargo manifest as mistargeting of apicoplast proteins to the PV (28). To assess whether stabilization of ACP_L_-GFP-mDHFR blocked protein import early during sorting or later after arrival at the apicoplast, we used live-and fixed-cell imaging to assess whether this fusion protein and the endogenous apicoplast protein ACP were successfully sorted in WR99210-treated parasites. As expected, in untreated parasites ACP_L_-GFP-mDHFR localized to an elongated structure characteristic of the apicoplast and co-localized with ACP (Fig. 4A and B). After 1 day of WR99210 treatment (i.e., within the same lytic cycle as treatment), the majority of cells exhibited ACP_L_-GFP-mDHFR and ACP signal in a single punctum or elongated pattern that likely indicates an intact apicoplast (Fig. 4A and B). After 3 days of WR99210 treatment (i.e., after 1 full lytic cycle of treatment), both ACP_L_-GFP-mDHFR and ACP were present almost exclusively in diffuse puncta that are thought to represent vesicles post-sorting in parasites that have lost their apicoplast (30) (Fig. 4A and B). Altogether, the localization of ACP_L_-GFP-mDHFR and ACP to either organelle-like structures or diffuse puncta following mDHFR stabilization indicates that sorting in the parasite endomembrane system remained intact. This suggests that the defect in WR99210-treated ACP_L_-GFP-mDHFR parasites occurs after this initial protein sorting step.

**Figure 4.**
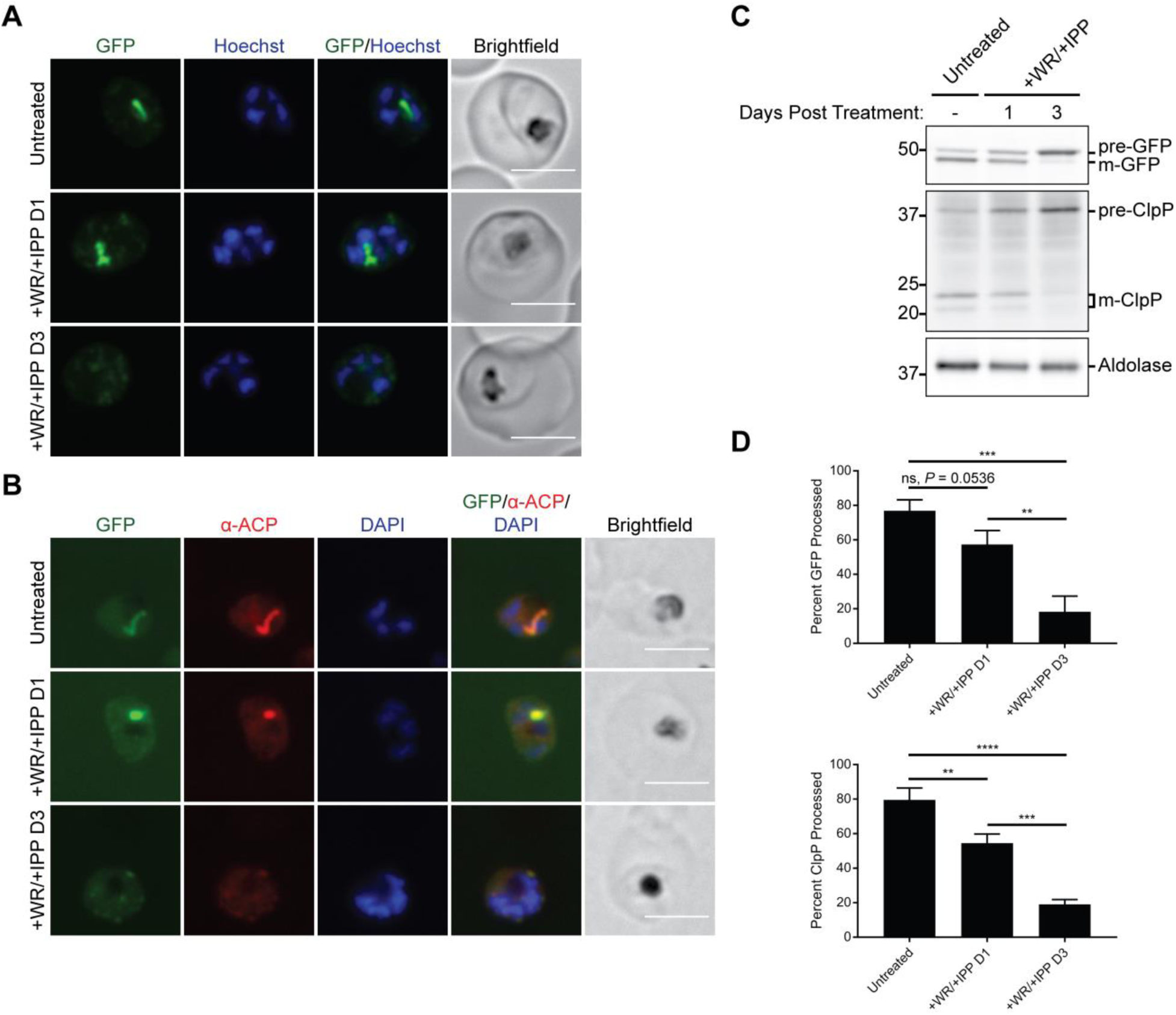
Conditional stabilization of apicoplast-targeted GFP-mDHFR disrupts import of endogenous apicoplast cargo across the apicoplast membranes. ACP_L_-GFP-mDHFR parasites were grown with 10 nM WR99210 and 200 µM IPP for either 1 or 3 days (corresponding to the same lytic cycle or the first lytic cycle post-treatment, respectively). (A) Live imaging of Hoechst-stained parasites. Scale bars, 5 µm. (B) Fixed imaging of parasites stained with an antibody against the endogenous apicoplast marker ACP (recognizing an epitope not present on the ACP_L_ fused to GFP-mDHFR). Scale bars, 5 µm. (C) Western blot to assess transit peptide processing of ACP_L_-GFP-mDHFR and the endogenous apicoplast protein ClpP. pre-, precursor (unprocessed) protein; m-, mature (processed) protein. (D) Quantification of transit peptide processing for ACP_L_-GFP-mDHFR and ClpP. Data are expressed as the percentage of total GFP or ClpP signal that is mature (processed). Error bars represent standard deviation of the mean of 3 biological replicates. \*\**P* < 0.01, \*\*\**P* < 0.001, \*\*\*\**P* < 0.0001, one-way ANOVA with Tukey’s multiple comparisons test. ns, not significant.

Most apicoplast-targeted proteins contain an *N*-terminal transit peptide that is proteolytically processed upon successful import into the organelle lumen, and an accumulation of unprocessed protein can be used as a marker for a protein import defect. We expected that stabilized ACP_L_-GFP-mDHFR would block translocation of apicoplast cargo into the organelle lumen and cause a transit peptide processing defect in WR99210-treated parasites. We therefore assessed the processing of the ACP_L_-GFP-mDHFR fusion and the endogenous apicoplast protein ClpP by western blotting. Consistent with a defect in protein translocation, WR99210-treated parasites exhibited a modest but reproducible accumulation of unprocessed ACP_L_-GFP-mDHFR after 1 day of WR99210 treatment (Fig. 4C and D). ClpP showed a comparable transit peptide processing defect (Fig. 4C and D), suggesting that stabilization of ACP_L_-GFP-mDHFR affected import not only of the fusion protein itself but also of endogenous apicoplast cargo. Processing of both the GFP-mDHFR fusion and ClpP was almost completely ablated after 3 days of WR99210 treatment (Fig. 4C and D). Because apicoplast proteins are still sorted to the apicoplast (Fig. 4A and B) but show a transit peptide processing defect (Fig. 4C and D), these data are consistent with a disruption in the translocation of apicoplast proteins across one or more apicoplast membranes.

In lieu of direct inhibitors of apicoplast protein import, we used a chemical biology approach to conditionally block apicoplast protein import by addition of a small molecule. Altogether, our results suggest a model in which chemically stabilized ACP_L_-GFP-mDHFR traffics to the apicoplast and stalls within putative membrane translocons, preventing import of endogenous apicoplast cargo and disrupting apicoplast biogenesis and function. Given that apicoplast protein import is likely required for biogenesis pathways such as apicoplast genome replication, the emergence of both a transit peptide processing defect (a readout for protein import; Fig. 4C and D) and a genome replication defect (Fig. 3) after 1 day of ACP_L_-GFP-mDHFR stabilization are consistent with this model. Unfortunately, because we could not detect direct biochemical interaction of stabilized ACP_L_-GFP-mDHFR with apicoplast translocons, we cannot rule out translocation-independent mechanisms-of-action, such as toxicity due to accumulation of stably folded ACP_L_-GFP-mDHFR in the apicoplast lumen. However, given the previous uses of mDHFR fusions to block protein translocation (24-27) and the defect in apicoplast protein import observed after just 1 day of WR99210 treatment (Fig. 4C and D), our model seems the most parsimonious.

These findings suggest that the proteins required for apicoplast protein import can serve as antimalarial targets that avoid delayed-death growth inhibition. Over a dozen proteins have been implicated in import of nuclear-encoded apicoplast proteins, but of particular interest are proteins related to established drug targets in other systems. For example, the apicoplast-localized AAA ATPase CDC48 is related to mammalian p97 and is likely involved in translocation of apicoplast cargo across the periplastid membrane (12). Notably, mammalian p97, which plays an important role in ERAD and other cellular processes, is of interest as an anti-cancer drug target. Specific inhibitors of mammalian p97 have been developed (31-34), suggesting that the same could be accomplished for the apicoplast-localized CDC48 in apicomplexans. Similarly, components of an apicoplast-localized ubiquitylation system are essential for apicoplast protein import and may also represent potential drug targets (11). A small molecule inhibitor of UAE, a ubiquitin-activating enzyme in humans, has recently been reported to have activity in multiple tumor models (35), indicating that the apicoplast-localized ubiquitin activating enzyme may also be a valuable drug target. The druggability of other members of apicoplast protein import complexes has not been explored and may yield additional targets.

Overall, our data suggest that chemical inhibition of apicoplast protein import machinery may be a viable strategy for development of next-generation antimalarials. Target-based drug discovery efforts against known import machinery may therefore yield specific inhibitors of apicoplast biogenesis with mechanisms-of-action orthogonal to those of current antimalarials.

## Materials and Methods

### Ethics statement

Human erythrocytes were purchased from the Stanford Blood Center (Stanford, California) to support in vitro *P*. *falciparum* cultures. Because erythrocytes were collected from anonymized donors with no access to identifying information, IRB approval was not required. All consent to participate in research was collected by the Stanford Blood Center.

### Parasite growth

*P. falciparum* Dd2^attB^ parasites (MRA-843) were obtained from MR4 and were grown in human erythrocytes (2% hematocrit) obtained from the Stanford Blood Center in RPMI 1640 media (Gibco) supplemented with 0.25% AlbuMAX II (Gibco), 2 g/L sodium bicarbonate, 0.1 mM hypoxanthine (Sigma), 25 mM HEPES, pH 7.4 (Sigma), and 50 μg/L gentamicin (Gold Biotechnology) at 37°C, 5% O2, and 5% CO2.

### Vector construction

Oligonucleotides (Table 1) were purchased from the Stanford Protein and Nucleic Acid facility and molecular cloning was performed using In-Fusion cloning (Clontech). GFP and mDHFR were PCR amplified from pARL2-SBP1-mDHFR-GFP (27) using primers MB119/MB120 and MB121/MB122, respectively. These products were simultaneously cloned into the BsiWi/AflII sites of the plasmid pRL2-ACP_L_-GFP (36) or a similar plasmid containing the ACP_L_ K18E mutant (AAA to GAA codon change) to generate pRL2-ACP_L_-GFP-mDHFR and pRL2-ACPL(K18E)-GFP-mDHFR for expression of GFP-mDHFR fusions from the mitochondrial ribosomal protein L2 promoter (37).

**Table 1.**
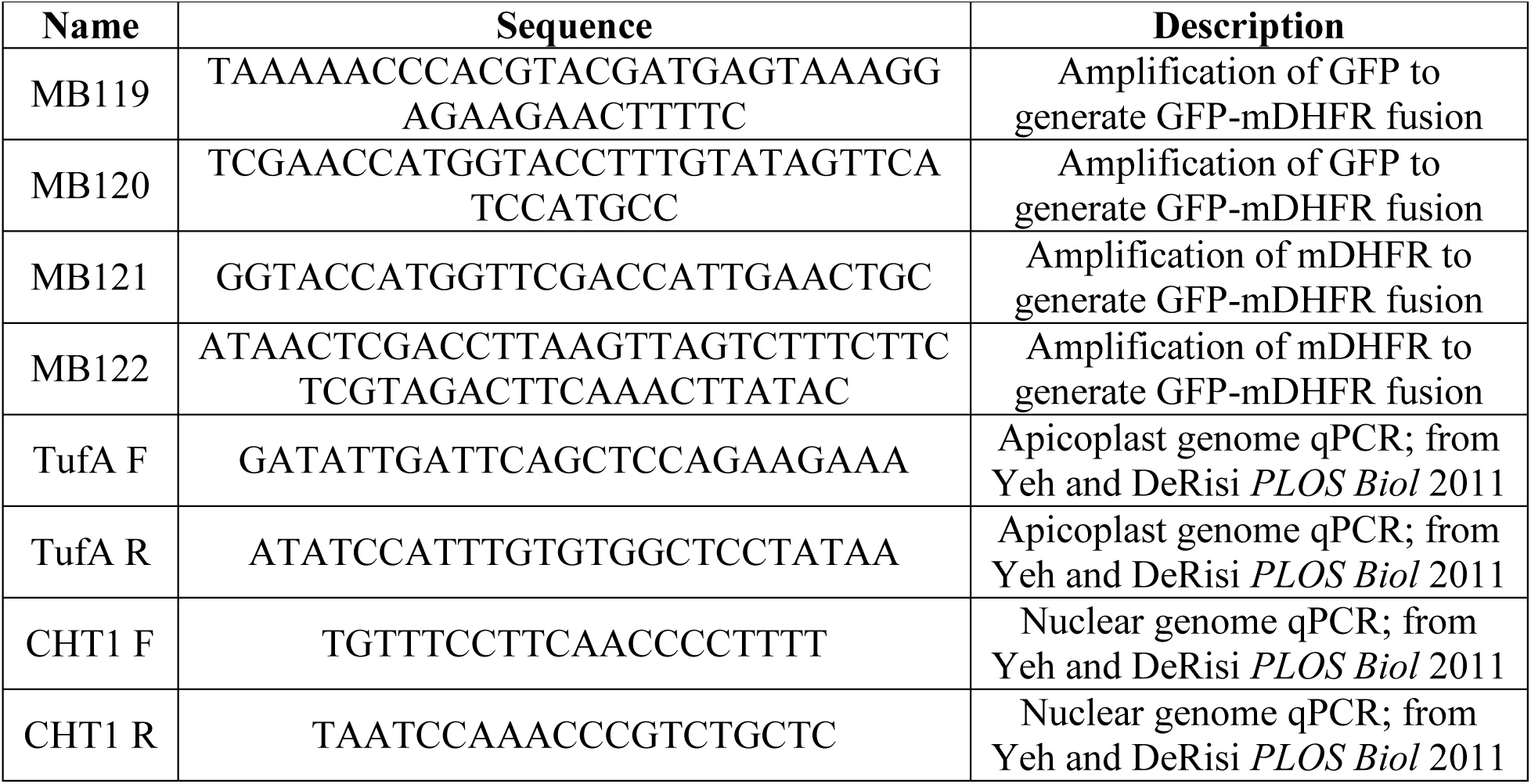
Primers used in this study.

### Parasite transfection

Transfections into Dd2^attB^ parasites were performed using a variation of the spontaneous uptake method (38, 39). Briefly, 50 μg each of pINT (29) and the desired pRL2 plasmid were ethanol precipitated and resuspended in a 0.2 cm electroporation cuvette in 100 μL TE buffer, 100 μL RPMI 1640 containing 10 mM HEPES-NaOH, pH 7.4, and 200 μL packed uninfected erythrocytes. Erythrocytes were pulsed with 8 square wave pulses of 365 V x 1 ms separated by 0.1 s and were allowed to reseal for 1 hour in a 37°C water bath before allowing parasites to invade. Drug selection with 2.5 μg/mL Blasticidin S (Research Products International) was initiated 4 days after transfection.

### Microscopy

For live imaging, parasites were settled onto glass-bottomed microwell dishes (MatTek P35G-1.5-14-C) in PBS containing 0.4% glucose and 2 μg/mL Hoechst 33342 stain (ThermoFisher H3570).

For fixed imaging, parasites were processed as previously described (40) with modifications. Briefly, parasites were washed in PBS and fixed in 4% paraformaldehyde (Electron Microscopy Sciences 15710) and 0.0075% glutaraldehyde (Electron Microscopy Sciences 16019) in PBS for 20 minutes. Cells were washed once in PBS, resuspended in PBS, and allowed to settle onto poly-L-lysine-coated coverslips (Corning) for 1 hour. Coverslips were washed once with PBS, permeabilized in 0.1% Triton X-100 in PBS for 10 minutes, and washed twice more in PBS. Coverslips were treated with 0.1 mg/mL sodium borohydride in PBS for 10 minutes, washed once in PBS, and blocked in 5% BSA in PBS. Following blocking, parasites were stained with 1:500 rabbit-α-*Pf*ACP (41) diluted in 5% BSA in PBS overnight at 4°C. Coverslips were washed three times in PBS, incubated for 1 hour in 1:3000 donkey-α-rabbit 568 secondary antibody (ThermoFisher A10042) in 5% BSA in PBS, washed three times in PBS, mounted onto slides with ProLong Gold antifade reagent with DAPI (ThermoFisher P36935), and sealed with nail polish prior to imaging.

Cells were imaged with a 100X, 1.35 NA objective on an Olympus IX70 microscope with a DeltaVision system (Applied Precision) controlled with SoftWorx version 4.1.0 and equipped with a CoolSnap-HQ CCD camera (Photometrics). Brightness and contrast were adjusted in Fiji (ImageJ) for display purposes.

### Parasite growth assays

For dose-response assays, sorbitol-synchronized ring-stage parasites were grown in 96-well plates containing 2-fold serial dilutions of WR99210 (Jacobus Pharmaceutical Company). After 3 days of growth, parasites were fixed in 1% paraformaldehyde in PBS and were stained with 50 nM YOYO-1 Iodide (ThermoFisher Y3601). Parasitemia was analyzed on a BD Accuri C6 flow cytometer. Each biological replicate of dose-response assays was performed in technical triplicate.

For time course growth experiments, sorbitol-synchronized parasites were untreated or were grown with 10 nM WR99210 with or without 200 μM IPP (Isoprenoids, LLC) for 5 days. Cultures were treated identically in terms of media changes and splitting into fresh erythrocytes. Samples to assess growth were collected daily, fixed in 1% paraformaldehyde in PBS, and stored at 4°C until completion of the experiment. Samples were then stained with YOYO-1 and analyzed as above.

### qPCR

Samples for DNA isolation were harvested daily during growth time course experiments. Parasites were released from erythrocytes by treatment with 0.1% saponin, washed in PBS, and stored at −80°C until analysis. Total parasite DNA was isolated using the DNeasy Blood & Tissue kit (Qiagen). qPCR was performed using Power SYBR Green PCR Master Mix (Thermo Fisher) with 0.15 µM each CHT1 F and CHT1 R primers targeting the nuclear gene chitinase or TufA F and TufA R primers targeting the apicoplast gene elongation factor Tu (30). qPCR was performed on Applied Biosystems 7900HT or ViiA 7 Real-Time PCR systems with the following thermocycling conditions: initial denaturation 95°C/10 minutes; 35 cycles of 95°C/1 minute, 56°C/1 minute, 65°C/1 minute; final extension 65°C/10 minutes. Relative quantification was performed using the ΔΔCt method.

### Western blotting

Sorbitol-synchronized parasites were untreated or were treated with 10 nM WR99210 and 200 μM IPP for 1 or 3 days. Parasites were separated from RBCs by lysis in 0.1% saponin, washed in PBS, and were stored at −80°C until analysis. Parasite pellets were resuspended in PBS containing 1X NuPAGE LDS sample buffer with 50 mM DTT and were heated to 95°C for 10 minutes before separation on NuPAGE Bis-Tris gels and transfer to nitrocellulose. Membranes were blocked in 0.1% Hammarsten casein (Affymetrix) in 0.2X PBS with 0.01% sodium azide. Antibody incubations were performed in a 1:1 mixture of blocking buffer and TBST (Tris-buffered saline with Tween 20: 10 mM Tris, pH 8.0, 150 mM NaCl, 0.25 mM EDTA, 0.05% Tween 20. Blots were incubated with primary antibody at 4°C overnight at the following dilutions: 1:20,000 mouse-α-GFP JL-8 (Clontech 632381); 1:4000 rabbit-α-*Pf*ClpP (42); 1:20,000 rabbit-α-*Pf*Aldolase (Abcam ab207494). Blots were washed once in TBST and were incubated for 1 hour at room temperature in 1:10,000 dilutions of IRDye 800CW donkey-α-rabbit or IRDye 680LT goat-α-mouse secondary antibodies (LI-COR Biosciences). Blots were washed three times in TBST and once in PBS before imaging on a LI-COR Odyssey imager. Band intensities of precursor and mature protein were quantified using Image Studio Lite version 5.2 (LI-COR).

### Statistics

One-way ANOVAs with Tukey’s multiple comparisons tests were performed in GraphPad Prism version 7.04.

## Acknowledgements

Funding for this work was provided by National Institutes of Health grants K08 AI097239 and DP5 OD012119 (E.Y.), a Burroughs Wellcome Fund Career Award for Medical Scientists (E.Y.), the Chan Zuckerberg Biohub Investigator Program (E.Y.), and a William R. and Sara Hart Kimball Stanford Graduate Fellowship (M.J.B.).

We thank Tobias Spielmann for providing plasmids encoding mDHFR, Sean Prigge for α-*Pf*ACP antibody, Walid Houry for α-*Pf*ClpP antibody, and Jacobus Pharmaceutical Company for WR99210.

## References

1. World Health Organization. 2017. World Malaria Report 2017. World Health Organization, Geneva, Switzerland.

2. Dondorp AM, Nosten F, Yi P, Das D, Phyo AP, Tarning J, Lwin KM, Ariey F, Hanpithakpong W, Lee SJ, Ringwald P, Silamut K, Imwong M, Chotivanich K, Lim P, Herdman T, An SS, Yeung S, Singhasivanon P, Day NP, Lindegardh N, Socheat D, White NJ. 2009. Artemisinin resistance in *Plasmodium falciparum* malaria. N Engl J Med 361:455–67.

3. Ashley EA, Dhorda M, Fairhurst RM, Amaratunga C, Lim P, Suon S, Sreng S, Anderson JM, Mao S, Sam B, Sopha C, Chuor CM, Nguon C, Sovannaroth S, Pukrittayakamee S, Jittamala P, Chotivanich K, Chutasmit K, Suchatsoonthorn C, Runcharoen R, Hien TT, Thuy-Nhien NT, Thanh NV, Phu NH, Htut Y, Han KT, Aye KH, Mokuolu OA, Olaosebikan RR, Folaranmi OO, Mayxay M, Khanthavong M, Hongvanthong B, Newton PN, Onyamboko MA, Fanello CI, Tshefu AK, Mishra N, Valecha N, Phyo AP, Nosten F, Yi P, Tripura R, Borrmann S, Bashraheil M, Peshu J, Faiz MA, Ghose A, Hossain MA, Samad R, Rahman MR, Hasan MM, Islam A, Miotto O, Amato R, MacInnis B, Stalker J, Kwiatkowski DP, Bozdech Z, Jeeyapant A, Cheah PY, Sakulthaew T, Chalk J, Intharabut B, Silamut K, Lee SJ, Vihokhern B, Kunasol C, Imwong M, Tarning J, Taylor WJ, Yeung S, Woodrow CJ, Flegg JA, Das D, Smith J, Venkatesan M, Plowe CV, Stepniewska K, Guerin PJ, Dondorp AM, Day NP, White NJ, Tracking Resistance to Artemisinin Collaboration (TRAC). 2014. Spread of artemisinin resistance in *Plasmodium falciparum* malaria. N Engl J Med 371:411–23.

4. McFadden GI, Reith ME, Munholland J, Lang-Unnasch N. 1996. Plastid in human parasites. Nature 381:482.

5. Kohler S, Delwiche CF, Denny PW, Tilney LG, Webster P, Wilson RJ, Palmer JD, Roos DS. 1997. A plastid of probable green algal origin in Apicomplexan parasites. Science 275:1485–9.

6. Sheiner L, Vaidya AB, McFadden GI. 2013. The metabolic roles of the endosymbiotic organelles of *Toxoplasma* and *Plasmodium* spp. Curr Opin Microbiol 16:452–8.

7. van Dooren GG, Striepen B. 2013. The algal past and parasite present of the apicoplast. Annu Rev Microbiol 67:271–89.

8. Spork S, Hiss JA, Mandel K, Sommer M, Kooij TW, Chu T, Schneider G, Maier UG, Przyborski JM. 2009. An unusual ERAD-like complex is targeted to the apicoplast of *Plasmodium falciparum*. Eukaryot Cell 8:1134–45.

9. Kalanon M, Tonkin CJ, McFadden GI. 2009. Characterization of two putative protein translocation components in the apicoplast of *Plasmodium falciparum*. Eukaryot Cell 8:1146–54.

10. Agrawal S, van Dooren GG, Beatty WL, Striepen B. 2009. Genetic evidence that an endosymbiont-derived endoplasmic reticulum-associated protein degradation (ERAD) system functions in import of apicoplast proteins. J Biol Chem 284:33683–91.

11. Agrawal S, Chung DW, Ponts N, van Dooren GG, Prudhomme J, Brooks CF, Rodrigues EM, Tan JC, Ferdig MT, Striepen B, Le Roch KG. 2013. An apicoplast localized ubiquitylation system is required for the import of nuclear-encoded plastid proteins. PLOS Pathog 9:e1003426.

12. Fellows JD, Cipriano MJ, Agrawal S, Striepen B. 2017. A plastid protein that evolved from ubiquitin and is required for apicoplast protein import in *Toxoplasma gondii*. mBio 8:e00950–17.

13. van Dooren GG, Tomova C, Agrawal S, Humbel BM, Striepen B. 2008. *Toxoplasma gondii* Tic20 is essential for apicoplast protein import. Proc Natl Acad Sci U S A 105:13574–9.

14. Glaser S, van Dooren GG, Agrawal S, Brooks CF, McFadden GI, Striepen B, Higgins MK. 2012. Tic22 is an essential chaperone required for protein import into the apicoplast. J Biol Chem 287:39505–12.

15. Sheiner L, Fellows JD, Ovciarikova J, Brooks CF, Agrawal S, Holmes ZC, Bietz I, Flinner N, Heiny S, Mirus O, Przyborski JM, Striepen B. 2015. *Toxoplasma gondii* Toc75 functions in import of stromal but not peripheral apicoplast proteins. Traffic 16:1254–69.

16. Goodman CD, Su V, McFadden GI. 2007. The effects of anti-bacterials on the malaria parasite *Plasmodium falciparum*. Mol Biochem Parasitol 152:181–91.

17. Dahl EL, Rosenthal PJ. 2007. Multiple antibiotics exert delayed effects against the *Plasmodium falciparum* apicoplast. Antimicrob Agents Chemother 51:3485–90.

18. de Koning-Ward TF, Gilson PR, Crabb BS. 2015. Advances in molecular genetic systems in malaria. Nat Rev Microbiol 13:373–87.

19. Rottmann M, McNamara C, Yeung BK, Lee MC, Zou B, Russell B, Seitz P, Plouffe DM, Dharia NV, Tan J, Cohen SB, Spencer KR, Gonzalez-Paez GE, Lakshminarayana SB, Goh A, Suwanarusk R, Jegla T, Schmitt EK, Beck HP, Brun R, Nosten F, Renia L, Dartois V, Keller TH, Fidock DA, Winzeler EA, Diagana TT. 2010. Spiroindolones, a potent compound class for the treatment of malaria. Science 329:1175–80.

20. Ganesan SM, Falla A, Goldfless SJ, Nasamu AS, Niles JC. 2016. Synthetic RNA-protein modules integrated with native translation mechanisms to control gene expression in malaria parasites. Nat Commun 7:10727.

21. Amberg-Johnson K, Hari SB, Ganesan SM, Lorenzi HA, Sauer RT, Niles JC, Yeh E. 2017. Small molecule inhibition of apicomplexan FtsH1 disrupts plastid biogenesis in human pathogens. Elife 6:e29865.

22. Armstrong CM, Goldberg DE. 2007. An FKBP destabilization domain modulates protein levels in *Plasmodium falciparum*. Nat Methods 4:1007–9.

23. Muralidharan V, Oksman A, Iwamoto M, Wandless TJ, Goldberg DE. 2011. Asparagine repeat function in a *Plasmodium falciparum* protein assessed via a regulatable fluorescent affinity tag. Proc Natl Acad Sci U S A 108:4411–6.

24. Eilers M, Schatz G. 1986. Binding of a specific ligand inhibits import of a purified precursor protein into mitochondria. Nature 322:228–32.

25. Gehde N, Hinrichs C, Montilla I, Charpian S, Lingelbach K, Przyborski JM. 2009. Protein unfolding is an essential requirement for transport across the parasitophorous vacuolar membrane of *Plasmodium falciparum*. Mol Microbiol 71:613–28.

26. Gruring C, Heiber A, Kruse F, Flemming S, Franci G, Colombo SF, Fasana E, Schoeler H, Borgese N, Stunnenberg HG, Przyborski JM, Gilberger TW, Spielmann T. 2012. Uncovering common principles in protein export of malaria parasites. Cell Host Microbe 12:717–29.

27. Mesen-Ramirez P, Reinsch F, Blancke Soares A, Bergmann B, Ullrich AK, Tenzer S, Spielmann T. 2016. Stable translocation intermediates jam global protein export in *Plasmodium falciparum* parasites and link the PTEX component EXP2 with translocation activity. PLOS Pathog 12:e1005618.

28. Foth BJ, Ralph SA, Tonkin CJ, Struck NS, Fraunholz M, Roos DS, Cowman AF, McFadden GI. 2003. Dissecting apicoplast targeting in the malaria parasite *Plasmodium falciparum*. Science 299:705–8.

29. Nkrumah LJ, Muhle RA, Moura PA, Ghosh P, Hatfull GF, Jacobs WR, Jr., Fidock DA. 2006. Efficient site-specific integration in *Plasmodium falciparum* chromosomes mediated by mycobacteriophage Bxb1 integrase. Nat Methods 3:615–21.

30. Yeh E, DeRisi JL. 2011. Chemical rescue of malaria parasites lacking an apicoplast defines organelle function in blood-stage *Plasmodium falciparum*. PLOS Biol 9:e1001138.

31. Chou TF, Brown SJ, Minond D, Nordin BE, Li K, Jones AC, Chase P, Porubsky PR, Stoltz BM, Schoenen FJ, Patricelli MP, Hodder P, Rosen H, Deshaies RJ. 2011. Reversible inhibitor of p97, DBeQ, impairs both ubiquitin-dependent and autophagic protein clearance pathways. Proc Natl Acad Sci U S A 108:4834–9.

32. Chou TF, Li K, Frankowski KJ, Schoenen FJ, Deshaies RJ. 2013. Structure-activity relationship study reveals ML240 and ML241 as potent and selective inhibitors of p97 ATPase. ChemMedChem 8:297–312.

33. Magnaghi P, D’Alessio R, Valsasina B, Avanzi N, Rizzi S, Asa D, Gasparri F, Cozzi L, Cucchi U, Orrenius C, Polucci P, Ballinari D, Perrera C, Leone A, Cervi G, Casale E, Xiao Y, Wong C, Anderson DJ, Galvani A, Donati D, O’Brien T, Jackson PK, Isacchi A. 2013. Covalent and allosteric inhibitors of the ATPase VCP/p97 induce cancer cell death. Nat Chem Biol 9:548–56.

34. Anderson DJ, Le Moigne R, Djakovic S, Kumar B, Rice J, Wong S, Wang J, Yao B, Valle E, Kiss von Soly S, Madriaga A, Soriano F, Menon MK, Wu ZY, Kampmann M, Chen Y, Weissman JS, Aftab BT, Yakes FM, Shawver L, Zhou HJ, Wustrow D, Rolfe M. 2015. Targeting the AAA ATPase p97 as an approach to treat cancer through disruption of protein homeostasis. Cancer Cell 28:653–665.

35. Hyer ML, Milhollen MA, Ciavarri J, Fleming P, Traore T, Sappal D, Huck J, Shi J, Gavin J, Brownell J, Yang Y, Stringer B, Griffin R, Bruzzese F, Soucy T, Duffy J, Rabino C, Riceberg J, Hoar K, Lublinsky A, Menon S, Sintchak M, Bump N, Pulukuri SM, Langston S, Tirrell S, Kuranda M, Veiby P, Newcomb J, Li P, Wu JT, Powe J, Dick LR, Greenspan P, Galvin K, Manfredi M, Claiborne C, Amidon BS, Bence NF. 2018. A small-molecule inhibitor of the ubiquitin activating enzyme for cancer treatment. Nat Med 24:186–193.

36. Boucher MJ, Ghosh S, Zhang L, Lal A, Jang SW, Ju A, Zhang S, Wang X, Ralph SA, Zou J, Elias JE, Yeh E. 2018. Integrative proteomics and bioinformatic prediction enable a high-confidence apicoplast proteome in malaria parasites. PLOS Biol 16:e2005895.

37. >Balabaskaran Nina P, Morrisey JM, Ganesan SM, Ke H, Pershing AM, Mather MW, Vaidya AB. 2011. ATP synthase complex of *Plasmodium falciparum*: dimeric assembly in mitochondrial membranes and resistance to genetic disruption. J Biol Chem 286:41312–22.

38. Deitsch K, Driskill C, Wellems T. 2001. Transformation of malaria parasites by the spontaneous uptake and expression of DNA from human erythrocytes. Nucleic Acids Res 29:850–3.

39. Wagner JC, Platt RJ, Goldfless SJ, Zhang F, Niles JC. 2014. Efficient CRISPR-Cas9-mediated genome editing in *Plasmodium falciparum*. Nat Methods 11:915–8.

40. Tonkin CJ, van Dooren GG, Spurck TP, Struck NS, Good RT, Handman E, Cowman AF, McFadden GI. 2004. Localization of organellar proteins in *Plasmodium falciparum* using a novel set of transfection vectors and a new immunofluorescence fixation method. Mol Biochem Parasitol 137:13–21.

41. Gallagher JR, Prigge ST. 2010. *Plasmodium falciparum* acyl carrier protein crystal structures in disulfide-linked and reduced states and their prevalence during blood stage growth. Proteins 78:575–88.

42. El Bakkouri M, Pow A, Mulichak A, Cheung KL, Artz JD, Amani M, Fell S, de Koning-Ward TF, Goodman CD, McFadden GI, Ortega J, Hui R, Houry WA. 2010. The Clp chaperones and proteases of the human malaria parasite *Plasmodium falciparum*. J Mol Biol 404:456–77.

